# Hypermutability of integrated sequences of viral origin in a Chlorarachniophyte

**DOI:** 10.64898/2026.01.13.699255

**Authors:** Lisa Mettrop, Anna Lipzen, Gilles Mirambeau, Kerrie Barry, Igor V. Grigoriev, Gwenaël Piganeau, Marc Krasovec

## Abstract

Mutations provide the raw material for evolution, but mutation rates are not uniform across genomes. Using a mutation accumulation experiment in the marine phytoplankton *Bigelowiella natans*, we discovered extreme local variation in mutation rate: over 1000-fold differences across its nuclear genome. While the baseline single-nucleotide mutation rate is approximately 3.5×10^−10^ per site per generation, a common value for unicellular species, two genomic regions derived from integrated viruses exhibit strikingly elevated rates of about 6×10^−7^. These two regions show a distinctive mutational signature with almost exclusively T/A→C/G transitions, a pattern also found in other non-eukaryote derived sequences in *B. natans*, contrary to the usual GC to AT mutation bias. Notably, hypermutation occurs only on TpA dinucleotides, and only in a subset of experimental lines, suggesting a regulated process rather than random genomic instability. We propose that *B. natans* targets invading DNA through localized hypermutation, reminiscent of deamination-based antiviral defense systems in animals, prompting the idea of genome editing as a conserved immune system in eukaryotes.

**Significance Statement:** De novo mutations provide the raw material for adaptation, but at high frequencies they can compromise genome integrity. Here, we describe a hypermutable process targeting two integrated viral genomes in a chlorarachniophyte alga, resulting in a mutation rate 1000 times higher than in other regions and a very particular mutation spectrum. These observations are reminiscent of hypermutation-based antiviral defenses described in humans against HIV and influenza; whereby host-mediated deamination of the viral genome increases its mutation rate such that the virus loses its infectivity.

## Introduction

*De novo* mutations continually change the genome, sustaining the capacity for evolutionary innovation. The rate of spontaneous mutation (*μ*), that is the number of mutations per base per generation, and the mutation spectrum, the relative frequency of different types of mutations, define this new diversity. *μ* is a quantifiable parameter under the influence of both selection and drift, and has been shown to span four orders of magnitude of variation within the tree of life (1). The mutation rate also presents large intracellular variations. In particular, eukaryotic cells contain multiple genomic compartments such as nuclear and mitochondrial genomes, as well as plastid genomes in photosynthetic organisms. For each of these compartments, distinct *μ* have been estimated. For example, in studied metazoans and unicellular eukaryotes, *μ*_*mito*_ is usually estimated to be higher than *μ*_*nuclear*_ (2, 3) while in land plants *μ*_*mito*_ is significantly lower than *μ*_*nuclear*_ (4). In contrast, *μ*_*chloroplast*_ is usually lower than *μ*_*nuclear*_ in both multicellular and unicellular photosynthetic organisms (4). Moreover, within the nuclear compartment itself, different types of sequences or regions with specific *μ* have been observed. For example, transcription can change the spontaneous mutation rate of coding sequences (5) and local mutation rate may be influenced by DNA methylation (6), chromatin structure (7) or histone marks (8). Significant differences in spontaneous mutation rates have also been observed between chromosomes in *Vibrio* bacteria (9). Another interesting sub-compartment is formed by sequences of exogenous origin, such as DNA viruses, which are integrated in many unicellular eukaryotic genomes (10, 11). Normally, viruses have a rapidly evolving genome, with mutation rates of around 10^-6^ -10^-4^ mutations per base for RNA viruses and 10^-8^-10^-6^ for DNA viruses (12) while eukaryotic genome mutation rates mostly range from 10^-10^ to 10^-8^ mutations per base per replication (1). However, the replication of endogenized viruses being dependent on the host cell, it could be that local mutation rate of integrated sequences approaches the global mutation rate of the host genome. Here, we conducted a mutation accumulation (MA) experiment in *Bigelowiella natans*. In MA experiments, population sizes of MA lines are kept extremely small with frequent bottlenecks (13). In this way, effective population size (*N*_*e*_) stays sufficiently small that selection is negligeable; hence, mutations are fixed independently of their effect on fitness. *B. natans* is a chlorarachniophyte, unicellular marine alga in the supergroup Rhizaria. Their chloroplast comes from a secondary endosymbiosis with a green alga. Interestingly, remnants of the symbiont nuclear genome are preserved as a nucleomorph, forming a fourth sub-cellular genomic compartment located between the outer membrane pairs of the *B. natans* chloroplast (14). Moreover, the genome of *B. natans* contains numerous integrated sequences of viral origin, predominantly derived from nucleocytoplasmic large DNA viruses (NCLDV) and virophages (15) (Fig. S1). This makes *B. natans* an ideal model to study intracellular variation of mutation rate and spectrum, both between several genomic compartments and within the nuclear genome.

## Results

### Mutation accumulation (MA) experiment

The MA experiment using the *B. natans* strain RCC623 lasted a total of 378 days with 27 bottlenecks, resulting in 15 successfully sequenced MA lines by Illumina NovaSeq for a total of 3,132 accumulated generations. The callable nuclear genome (*G\**), meaning the genomic sites usable for the analysis by applying mapping quality and depth thresholds, is 85,048,595 nucleotides (nt), so 90% of the genome size (14). Callable genomes for organelles and nuclear sub-compartments are detailed in Tables 1 and S1; no de novo mutations were found in the organellar and nucleomorph genomes, which can be explained by their small *G\**. Regarding the nucleomorph, we redid an analysis omitting the 2 MA lines with the lowest coverage to increase its *G\** from 37% to 58%, however, this still did not yield any de novo mutations. In the nuclear genome, a total of 302 single nucleotide mutations (SNM) were identified, as well as 21 insertions-deletions (ID) (Table S2 and S3). The SNM presented an extreme heterogeneity in their distribution in the nuclear genome (Fig. 1, Table 1 and Fig. S1) Additionally, using raw read depth as a proxy for sequence copy number, we found three candidates for de novo duplications (Fig. S2): in MA line 19 there was a large duplication in the middle of scaffold 53 from approximately position 169,900 to 470,200 (300,300bp), in MA line 15 a part of scaffold 61 was duplicated from approximately position 258,900 to 591,300 (332,400bp), and in MA line 21 the whole scaffold 74 was duplicated (544,257bp).

**Table 1.**
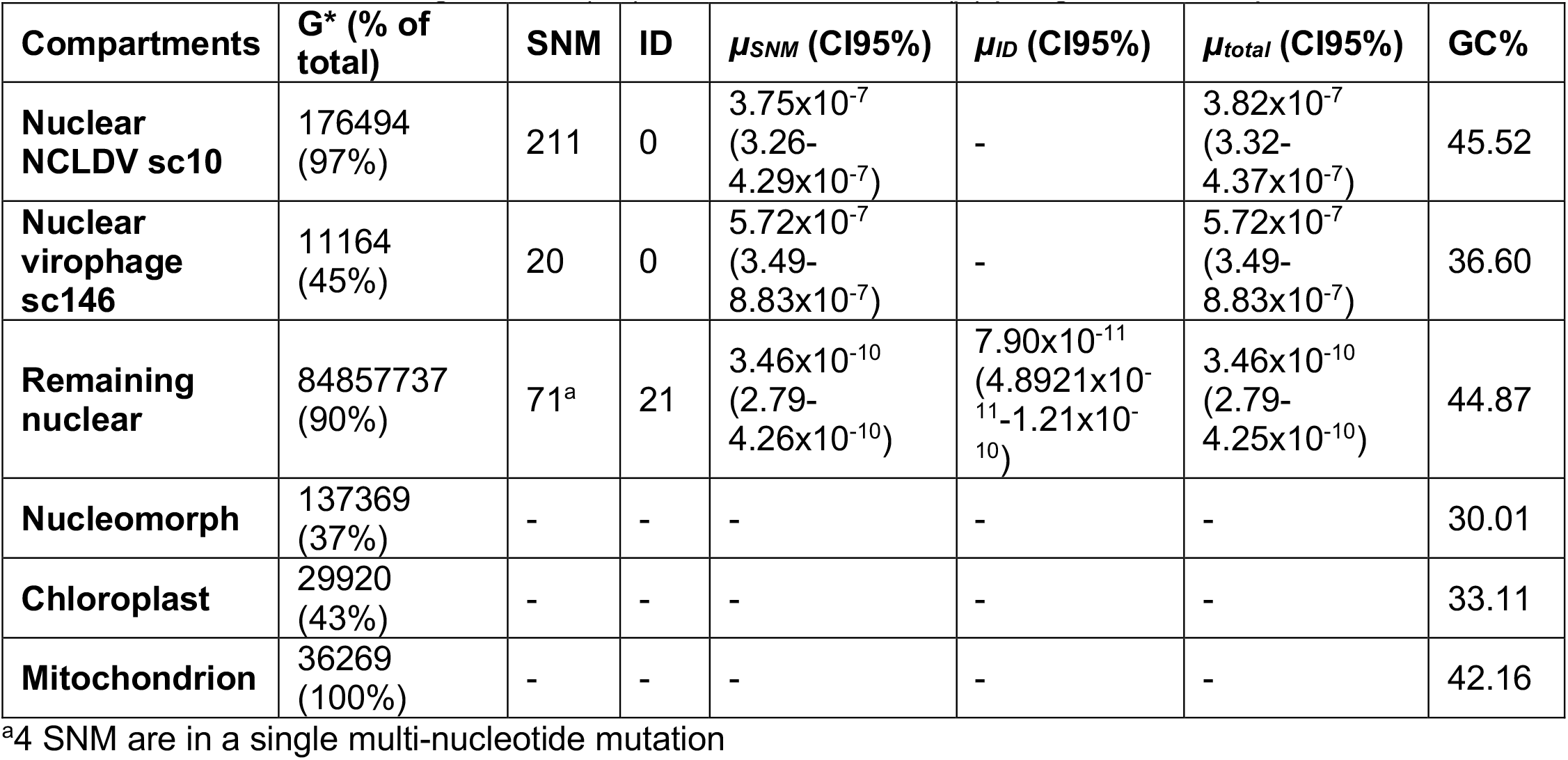
Size of the callable genomes (*G\**) and mutation rates (*μ*) per genomic compartment.

**Figure 1.**
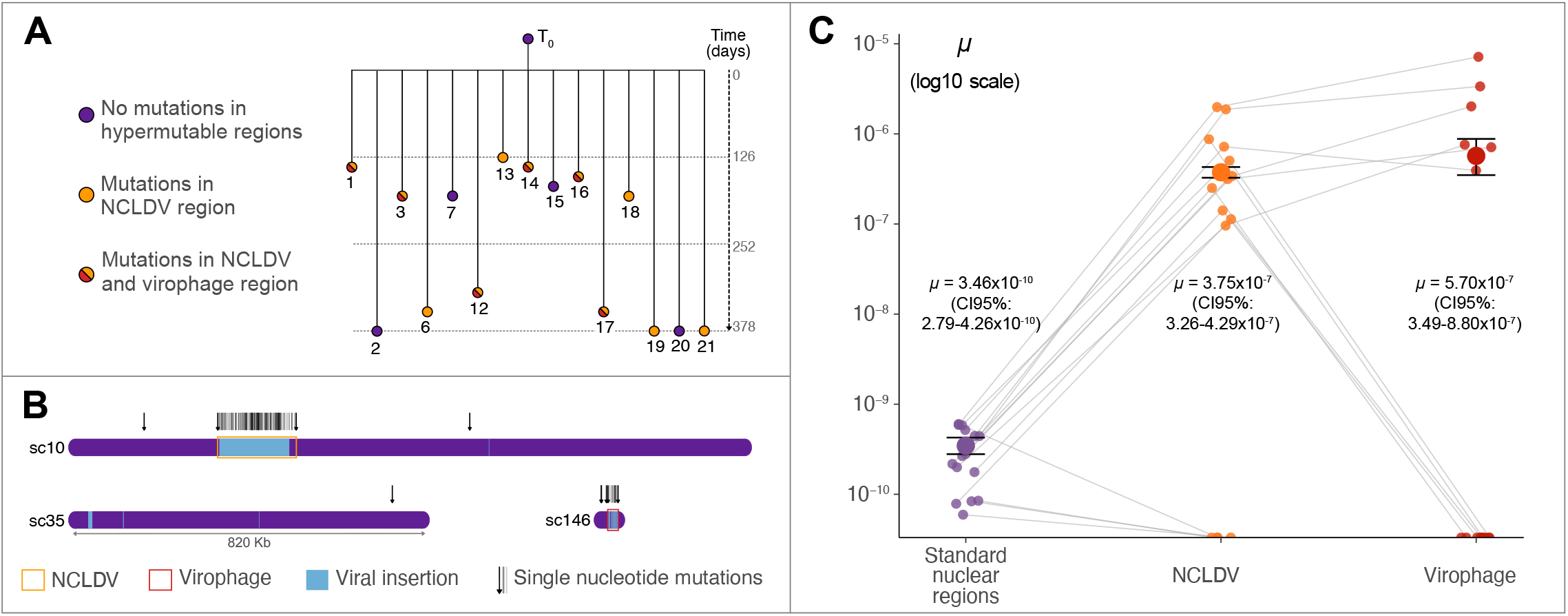
Organisation of the mutation accumulation (MA)-experiment, genome architecture, and mutation rate variability in *B. natans*. **A.** The MA experiment was started with a clonal culture (T_0_) from which MA lines were inoculated after single-cell isolation. The MA experiment lasted a total of 378 days for the MA lines that lasted the longest, with 15 MA lines that were usable for the mutation rate analysis. The numbers under the circles give the MA line identities. The colors of the circle of each MA line indicates if there were mutations in the NCLDV hypermutable region (orange), in the NCLDV and virophage hypermutable regions (red/orange) or in none of these (violet); there were no lines that had mutations in the virophage but not in the NCLDV hypermutable region. **B**. An example of the architecture of the *B. natans* genome, which contains many inserted ORFs of viral origins (NCLDV, virophage or transpoviron origins, indicated by blue blocks), as identified previously (15). Single nucleotide mutations are represented as black arrows (or bars in hypermutable regions) above the scaffold. Represented are scaffold 10, which contains a large hypermutable region of NCLDV origins (sc10:336000-521500) (orange rectangle); scaffold 35, which contains a viral insertion and presents a mutation, but has no hypermutable regions; and scaffold 146, which contains a hypermutable region of virophage origin (sc146:17000-41997) (red rectangle). **C**. The large variation of single nucleotide mutation rates between MA lines in the nuclear genome; the standard nuclear genome is the nuclear genome without the hypermutable regions of NCLDV and virophage origins. The average mutation rate *µ* with the 95% confidence interval for each compartment is written in the graph and represented by a large colored dot with horizontal bars for the confidence interval. The smaller translucent dots represent the mutation rates of each MA line, grey lines connect the values of a given MA line across compartments.

### Hypermutable regions of viral origin

There are two hypermutable regions in the nuclear genome that together contain 231 out of the 302 nuclear SNM. The larger hypermutable region is on scaffold 10, with 211 AT→GC SNM within positions 336,000-521,500 (±186 kb). This region has previously been reported to be of viral origin, notably an inserted NCLDV (15), although we extend it by ±21 kb here, based on the continuous high mutation rate. It stands out from other virus insertions in *B. natans* genome by its size (Fig. S3) and its transcriptional silence (15), see Fig. S4 for RNA sequencing depth from the MMETPS dataset (16). Using sequencing depth and Blastn, we confirmed that this NCLDV region is present in a single copy in the genome (Fig. S5). A second, smaller hypermutable region of virophage origin is present on the end of the scaffold 146 (positions 17,000-41,997, ±25 kb), with 20 exclusively AT→GC SNM. This region is densely packed with virus-like ORFs, and does not differ from other virophage insertions by size (Fig. S3) or gene expression (15). These two regions (hereafter called ‘viral hypermutable regions’) have extremely high mutation rates in the order of 10^-7^ mutation per site per generation, three orders of magnitude higher than the rest of the nuclear genome (Table 1, Table S4, Fisher’s exact test p-value < 2.2×10^-16^). Outside of these viral hypermutable regions, the nuclear single nucleotide mutation rate (*μ*_*SNM*_) is 3.46×10^-10^ and insertion-deletion rate *(μ*_*ID*_) is 7.90×10^-11^ (Table 1); these values being quite standard for unicellular eukaryotes (1), we will focus on the viral hypermutable regions in the rest of this paper. Interestingly, the hypermutation of viral regions was not observed in all MA lines (Fig. 1, Tables 2 and S3): of the 15 lines, 11 showed hyper-mutation of A and T residues on scaffold 10, ranging from 2 to 45 SNM in a single line (Fig. S6); in the line with most mutations, *μ*_*SNM*_ was as high as 1.98×10^-6^ in the NCLDV region (Table 2). On scaffold 146, only 6 lines had mutations in the viral region, and all of these also had mutations on scaffold 10. The heterogeneity of *μ*_*SNM*_ between lines suggests that the putative mutational mechanism at work is not always active.

**Table 2.** Number of de novo nucleotide mutations per MA line. *µ*_*SNM*_ viral considers all SNM in both NCLDV and virophage hypermutable regions with the combined *G\**_*viral*_ = 187,656 nt.

| Line | Generations | SNM nuclear | SNM NCLDV | SNM virophage | ID nuclear | $\mu_{SNM}$ viral |
| --- | --- | --- | --- | --- | --- | --- |
| 1 | 100 | 5 | 35 | 8 | 0 | $2.25 \times 10^{-6}$ |
| 12 | 252 | 6 | 14 | 2 | 2 | $3.33 \times 10^{-7}$ |
| 13 | 101 | 5 | 9 | 0 | 2 | $4.67 \times 10^{-7}$ |
| 14 | 118 | 2 | 2 | 1 | 0 | $1.33 \times 10^{-7}$ |
| 15 | 137 | 7 | 0 | 0 | 0 | 0 |
| 16 | 133 | 3 | 8 | 3 | 1 | $4.33 \times 10^{-7}$ |
| 18 | 150 | 1 | 3 | 0 | 0 | $1.05 \times 10^{-7}$ |
| 19 | 292 | 11 | 45 | 0 | 2 | $8.07 \times 10^{-7}$ |
| 2 | 278 | 2 | 0 | 0 | 0 | 0 |
| 20 | 397 | 2 | 0 | 0 | 0 | 0 |
| 21 | 401 | 6 | 10 | 0 | 3 | $1.31 \times 10^{-7}$ |
| 3 | 133 | 5 | 44 | 5 | 2 | $1.93 \times 10^{-6}$ |
| 17 | 228 | 10<sup>a</sup> | 29 | 1 | 4 | $6.89 \times 10^{-7}$ |
| 6 | 271 | 5 | 12 | 0 | 5 | $2.32 \times 10^{-7}$ |
| 7 | 141 | 1 | 0 | 0 | 0 | 0 |

<sup>a</sup>4 SNM are in a single multi-nucleotide mutation

### Striking change in mutation spectrum

In most known species, the mutation spectrum is AT-biased, meaning that G and C nucleotides mutate more frequently than A and T (17). While the spectrum of *B. natans* nuclear genome outside of the viral hypermutable regions is also AT-biased (*µ*_*GC>AT*_/*µ*_*AT>GC*_ = 2.65, Table S5), the viral hypermutable regions present an extreme bias towards GC (Fig. 2). In the virophage region, the 20 SNM are exclusively A→G and T→C; In the NCLDV region, 210 out of 211 SNM are A→G or T→C, giving a mutation bias of *µ*_*GC>AT*_/*µ*_*AT>GC*_ = 0.006 (Table S5). Interestingly, all of these GC-biased SNM in the viral hypermutable regions fall on TpA dinucleotides (Table S2). In the case of A→G transitions, the mutated A is preceded by a T (TpA→TpG). The T→C transitions are the reverse complement of this, with each mutated T followed by an A (TpA→CpA).

**Figure 2.**
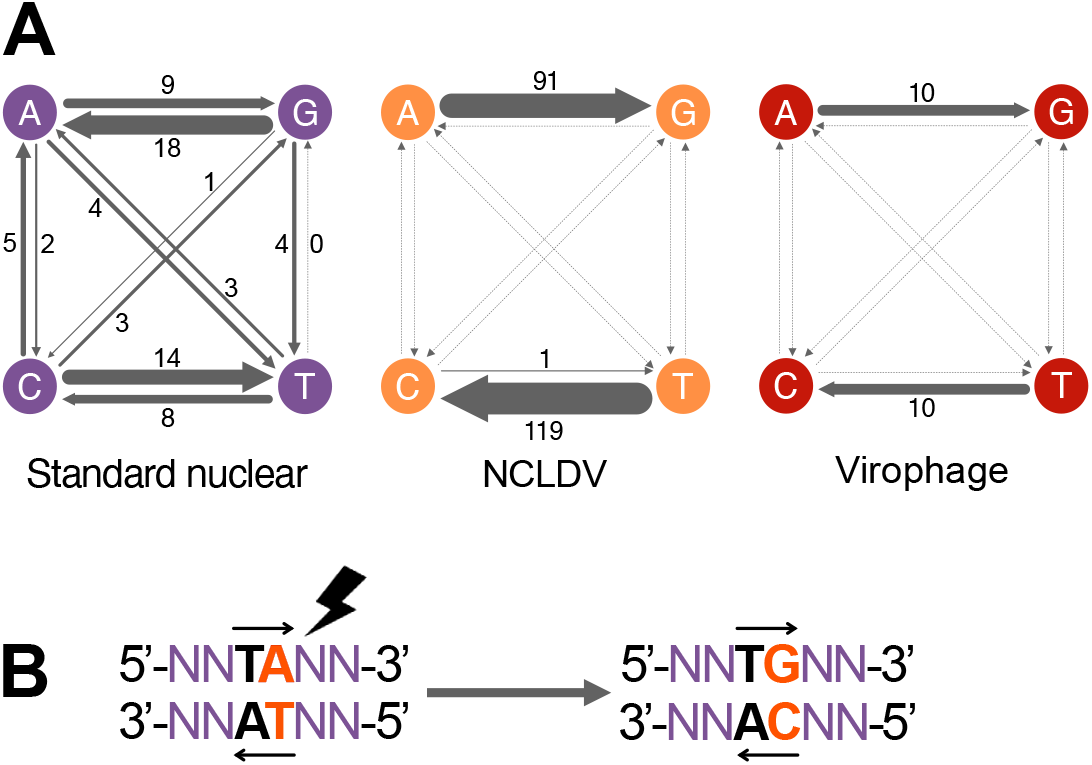
Nucleotide mutation spectrum in three different regions of the nuclear genome of *Bigelowiella natans*, and the specific mutation motif in the hypermutable regions. **A.** While the standard nuclear genome presents all 4 types of transitions plus 7 transversions (*μ*_*AT*→*GC*_ = 1.30×10^-10^ and *μ*_*GC*→*AT*_ = 3.44×10^-10^, thus AT-bias = 2.65, Ts/Tv = 7), mutation in the viral hypermutable regions are almost exclusively A→G and T→C transitions, with *μ*_*AT*→*GC*_ = 6.85×10^-7^ and *μ*_*GC*→*AT*_ = 3.90×10^-9^ in the NCLDV region and *μ*_*AT*→*GC*_ = 9.02×10^-7^ and *μ*_*GC*→*AT*_ = 0 in the virophage region. **B**. The mutations in the hyperamutable viral regions almost exclusively fall on TpA dinucleotides, which mutate to TpG and CpA dinucleotides, depending on the strand.

### Dinucleotide frequencies in B. natans predicted coding sequences (CDS)

To test whether the TpA-centered mutation spectrum observed in viral hypermutable regions reflects an evolutionarily active targeted mutational mechanism, we searched for its predicted long-term genomic signature, namely TpA depletion coupled with CpA+TpG enrichment, in coding sequences to see if other sequences of foreign origin have already undergone hypermutation. We used three complementary approaches:

First, we looked at dinucleotide frequencies in the CDS of 197 previously identified genes in *B. natans* that have transpoviron, virophage or NCLDV origins (15). The mean relative frequency of CpA+TpG dinucleotides in these CDS was 0.153, significantly higher than the mean frequency of 0.137 for all nuclear CDS (Wilcoxon rank sum test, p-value 4.17×10^-8^). Unexpectedly, the mean relative TpA frequency in the 197 CDS was 0.052, also significantly higher than the mean frequency of 0.041 in all genes (Wilcoxon rank sum test, p-value 5.39×10^-5^). Because this set of foreign-derived genes was relatively small and did not yield a clear depletion signature, we continued with an unbiased coding genome-wide strategy.

Second, we looked at all predicted CDS in *B. natans* (see Methods). By plotting the distribution of TpA dinucleotides using the relative frequency of each dinucleotide, we identified a subgroup depleted in TpA, with an average TpA-frequency of 0.0018 (Fig. S7). There was no clear subgroup enriched in CpA+TpG. This low-TpA subset contained 324 predicted CDS (Table S6). Pfam annotation identified 245 distinct domains in 115 of these sequences. To determine whether specific functions were overrepresented, we compared Pfam frequencies in this subset against their frequencies across all 21,708 predicted CDS using Fisher’s exact test with Benjamini– Hochberg correction. Four Pfam domains were significantly enriched in TpA-depleted sequences (Table S7): three are consistent with a viral or transposable element ancestry (DDE-superfamily endonuclease, Endonuclease-reverse transcriptase, and Reverse transcriptase), and one is a zinc-finger motif.

Third, we repeated this analysis with four adjustments (see Methods). Using 5%-extreme cut-off values, we extracted the CDS that were both TpA-depleted and CpA+TpG-enriched (Table S8). This time, 56 sequences had Pfam hits, for a total of 119 different Pfams. 49 Pfams had significant adjusted p-values, with enrichments ranging from 0 to 53.74 (Table S9). Of the Pfams with an enrichment higher than 0, 11 suggested foreign origins, 24 were of likely host origin and 11 others were ambiguous. Three of the four previously found Pfams were recovered. Notably, the most strongly enriched Pfams were predominantly of foreign origin: among the 17 significant Pfams with >25-fold enrichment, 8 were almost certainly derived from viral, transposon, bacterial, or fungal sources.

### Dinucleotide frequencies at whole genome scale

We then extended our search for sequences with low TpA and high CpA+TpG to the whole genome, which was cut into 1kb windows. For each window, a Rho value was calculated with the nucleotide composition to predict expected dinucleotide frequencies: Rho<1 means a depletion, Rho>1 means an enrichment (18). Plotting Rho_TpA_ versus the Rho of other dinucleotides shows that for a subgroup of windows, Rho_CpA+TpG_ increases strongly as Rho_TpA_ decreases in a linear fashion. To increase specificity, we re-calculated Pearson regression coefficients for all windows with Rho_TpA_ <0.5 and indeed, in this case, the negative relationship between Rho_TpA_ and Rho_CpA+TpG_ is the strongest (Rho=-0.5, p=0), suggesting that for genomic regions depleted in TpA, CpA+TpG increases in a linear fashion (Fig. S8).

Then, we tested whether the TpA-depleted/CpA+TpG-enriched windows form clusters. Again, we used 5% cut-off values to define these windows (Rho_TpA_ ≤ 0.43 and Rho_CpA+TpG_ ≥ 1.52, Fig. S9). Clusters were defined as at least three subsequent TpA-depleted/CpA+TpG-enriched windows (Figs. S10 and S11, Table S10). To evaluate whether the number of clusters was greater than expected by chance, we performed 10,000 permutations, shuffling Rho values within scaffolds. No permuted dataset produced as many clusters as observed (max 3 clusters versus 124 observed clusters, permutation p-value <0.0001). Interestingly, the whole scaffold 226 is a cluster (5 windows, 4360 bp). The 124 observed clusters contain 96 genes of which 18 have IPR annotations: 4 are viral and transposon related; 2 others possibly have bacterial origins (Table S11). The remaining 78 proteins had no annotation, including the one on scaffold 226. Clusters contain a significantly higher proportion of non-annotated proteins than the general genome (81.23% versus 57.94%, Fisher’s exact test p-value = 2.7×10^-5^).

Finally, we compared TpA frequency of the complete hypermutable NCLDV region to that of related giant virus genomes. Blasting the NCLDV sequence against the NCBI database shows that the region is probably of Phycodnaviridae provenance, as reported in (15) for the major capsid protein. Thus, we retrieved all complete Phycodnaviridae genomes from NCBI (n=38 on 4 March 2026, Fig. S12) and calculated Rho_TpA_ as described above. The NCLDV genome exhibited the lowest Rho_TpA_ value (Rho_TpA_ = 0.45) among all genomes examined and fell 2.62 standard deviations below the mean of the Phycodnaviridae distribution (Rho_TpA_ = 0.75 ± 0.13) (empirical test, p-value = 0.026).

### Signatures of long-term hypermutation in the T_0_ genome

To detect if the hypermutability of the viral hypermutable regions we observed in our MA experiment reflects a long-term hypermutability, we analyzed the neutral genetic diversity in the NCLDV region between the reference *B. natans* genome sequenced in 2012 (14) and our *T*_*0*_ sampled in 2021. Only single nucleotide polymorphisms (SNPs) in intergenic regions were considered. 158 SNPs were located in the NCLDV region on scaffold 10, resulting in a high diversity of 4.23×10^-3^ SNP per nucleotide (Table S12). For the remaining nuclear genome this was 1.54^-4^ with 4,419 SNPs, 27 times lower than on the NCLDV region (Fisher’s exact test, p-value < 2.2×10^-16^). On the virophage region, no SNPs fell in intergenic regions.

We also evaluated the SNP spectrum, assuming the alleles in the reference genome are the ancestral version of the variants found in our *T*_*0*_. We counted 144 intergenic SNPs from A/T→G/C in the NCLDV region and 14 SNPs from G/C→A/T. Considering sequence composition, this gives an AT-bias of 0.14, meaning A/T basepairs mutate 7.3 times faster to G/C basepairs than the inverse in this region (Table S12). In nuclear regions outside the viral regions, the AT-bias is 1.32, which is 9.4 times higher than in the NCLDV region (Fisher’s exact test, p-value < 2.2×10^-16^). However, even with this strong mutational bias, TpA mutational targets persist, since, in the reference genome, 3.3% (N = 6169) of dinucleotides in the NCLDV region are TpA, which is 1.8 times lower than the relative TpA frequency in the nuclear genome outside the viral regions (6.0%, (two-sample proportions test, p-value < 2.2×10^-16^). Interestingly, relative TpA frequency in the virophage region was 9.2% (N = 2300), higher than in the remaining nuclear genome (two-sample proportions test, p-value < 2.2×10^-16^).

### Adenosine (A)-deaminase candidates in B. natans

One explanation for the observed high AT→GC SNM rate in the hypermutable viral regions could be the activity of a DNA-editing mechanism, such as an A-deaminase. Adenosine can be deaminated into inosine, which is recognized as guanine during the next replication round (19). A-deaminases working on polynucleotides are universally conserved and involved in tRNA editing (TadA in bacteria, ADAT in eukaryotes) (20). Although it is classically assumed that ADATs edit RNA only, in amphioxus, BjADAT2 has recently been shown to edit DNA as well on A and C bases (21). Additionally, TbADAT2 in Trypanasoma can deaminate A in the RNA and C in the DNA (22). Thus, we searched for a DNA A-deaminase candidate in *B. natans*. Only one of the predicted *B. natans* proteins (Bn78255) possessed the conserved domains described for BjADAT2 and TbADAT2 (21) (File S1), most importantly the HxE-PCxxC motif and an N at position 119 (N113 in BjADAT2). This N seems pivotal for DNA editing capacity, as shown experimentally in *E. coli* EcTadA A-deaminase (23). We aligned the structural AlphaFold prediction of Bn78255 against the predicted structures of TbADAT2 and BjADAT2 in Pymol and found average RMSDs of 0.475 and 0.544 Å, respectively, meaning Bn78225 protein structure is highly similar to that of TbADAT2 and BjADAT2 (Fig. S13). Since BjADAT2 works together with BjADAT3 for DNA editing in the nucleus (21), we looked for a homologue of BjADAT3 and found a weaker candidate (Bn77986, 37.4% protein sequence identity over 155 amino acids). Although we cannot conclude on the activity of these proteins from predictions only, the presence of the ADAT2 homologue in *B. natans* makes hypermutation by A-deamination a reasonable hypothesis.

## Discussion

### Hypermutation as a defense against foreign sequences?

The extreme mutation rate and GC-mutation-bias in the viral hypermutable regions, very different from the rest of the nuclear genome, indicate that specific mutational processes may be at work. Since the hypermutable sequences are of viral origin, this hypermutation may be a host defense mechanism against invading sequences, comparable to repeat-induced point-mutation (RIP) in fungi, which targets sequences with high homology and causes strong C→T mutations, thus limiting the accumulation of transposable elements and other repeated sequences (24). Indeed, Pfams associated with predicted CDS that are TpA-depleted and CpA+TpG-enriched more often suggest a foreign origin than Pfams from other B. natans CDS, although many correspond to common eukaryotic domains. Three elements must be considered when interpreting these results. First, many TpA-depleted/CpA+TpG-enriched genes lack annotation, likely because they are already highly degraded. Second, transposons and integrated viruses are notoriously hard to annotate and may therefore be underrepresented in annotated sequences (25, 26). Third, some domains, like zinc-fingers, are intrinsically rich in TA, which is why we targeted sequences that are both TpA-depleted and CpA+TpG enriched. Altogether, our results suggest that hypermutation activity extends to other foreign sequences beyond integrated viruses. It is interesting that both virophage and NCLDV regions mutate at roughly the same rate, since these viruses play different roles during infection. NCLDV are known to prey on protists (27) and play an important role in biogeochemical cycles by lysis of eukaryotic phytoplankton (28). Virophages, on the other hand, are miniscule viruses that hijack NCLDV machinery (29) and have been proposed to act as a host defense system against NCLDV (15). Here, the putative hypermutation mechanism does not make any distinction between NCLDV and virophage.

### Inconsistent hypermutation of viral sequences

Hypermutation of viral sequences was not observed in all MA lines, which suggests that the mechanism is not constitutive. Then, why did it activate in some lines? There were no signs of viral infection during the MA experiment. However, a threat could come from the genome itself through the activation of transposable elements or lysogenic viruses, that integrate into the host genome and persist latently. Lysogenic viruses in unicellular eukaryotes have sparked interest recently; especially virophages are suspected of having lysogenic lifecycles (15, 29) but lysogenic NCLDV have also been described (30–32). Activity of such sequences may induce the putative defense mechanism, which could be guided through RNA intermediates, akin to known mechanisms like PIWI in animals (33) or RNA-directed-DNA-methylation in plants (34). Additionally, epigenetic modifications, including histone marks and DNA methylation, may play a role in the virus life cycle, such as for herpesviruses in humans (35). Since epimutation rate can be more than 10,000 times higher than genic mutation rate (36), it is possible that the hypermutation observed here was triggered by epigenetic mutations in multiple MA lines during the MA experiment.

### Genome editing, an immune system shared in eukaryotes?

The targeted mutation of TpA dinucleotides in DNA could correspond to an adenosine-deaminase activity which converts adenosine to inosine, thus causing A→G mutations. In line with this idea, DNA A-deamination has been demonstrated in the cell nucleus in Amphioxus (21), and in Trypanosoma ADAT-mediated C-deamination of the DNA is hypothesized to play a role in immunity by creating antigenic variation (22).

ADAT is related to immune systems in animals: ADAR proteins, thought to have evolved from ancestral ADAT after the acquisition of a double-stranded RNA binding domain (37), have been shown to act on RNA sequences of influenza and measles viruses as a defense in humans (38). ADAR has been well described in many metazoan lineages where it mostly targets repeat-derived dsRNA, and usually favours an U upstream of the mutated A (37), which is consistent with the TA-motif observed in this study. Additionally, in metazoans, the conserved AID/APOBEC proteins are also involved in immunity through deamination. They form a diverse group of enzymes that deaminate cytosines in both RNA and DNA substrates, leading to C→T mutation (39). The discovery of APOBEC was a key step in understanding cellular immunity to viruses, identifying hypermutation by DNA deamination as a mechanism inhibiting HIV replication (40).

APOBEC-like genes have been found beyond metazoans, notably in dinoflagellates, green algae and haptophytes (41). The origin of APOBEC in eukaryotes has been proposed to come from bacterial toxin systems, operating as mutators of hyper-variable genes, viruses and selfish elements (20). The parallels among ADAT-associated DNA editing in Amphioxus and Trypanosoma, ADAR/APOBEC-mediated defense against viral and repetitive sequences, and the potential hyperdeamination of foreign DNA in *B. natans* may reflect either convergent evolution of nucleic-acid-targeting deaminase activities or deep evolutionary conservation of an immune gene-editing function. The latter scenario is consistent with the recently proposed concept of ancestral immunity, which posits that many eukaryotic immune modules originate from prokaryotic systems (42). These modules are immune genes and domains, such as viperins and gasdermins, that are conserved across branches of eukaryotes and prokaryotes. Thus, although often considered clade-specific, the ancestral immunity hypothesis argues for a common and very ancient origin of immune proteins, in particular immunity against viruses, which probably exist since the first cell (43, 44). The ancestral immunity theory may explain the recurrence of similar defense systems across the tree of life and provides a compelling rationale for the resemblance between genome editing defenses in metazoans and *B. natans*.

## Conclusion

Here, we report a hypermutation of sequences of viral origin integrated in the genome of the chlorarachniophyte *B. natans*, with a very specific mutation spectrum. Based on these observations, we propose an often-overlooked role of mutation mechanisms: the protection of genome integrity against foreign sequences. These exciting prospects need to be studied further but suggest the existence of a specialized TpA hyper-deamination pathway in *B. natans* as a defense against integrating viruses and potentially transposable elements. Although this first study is not conclusive, the similarity of the observations in *B. natans* and systems discovered more than 20 years ago in humans by studying HIV and influenza genome hypermutability could point to genome-editing as a conserved anti-viral immune function.

## Materials and Methods

### Mutation accumulation (MA) experiment

*Bigelowiella natans* strain RCC623 (synonym to CCMP621) came from the Roscoff culture collection (45); its identity was confirmed by 18S rDNA PCR coupled with Sanger sequencing prior to the experiment. Cells were cultivated in L1 medium (Bigelow L1 Culture Medium Kit) generated from sterile natural sea water with salinity of 33 g/L in a light:dark cycle of 12:12 hours at 20°C. The mutation accumulation (MA) was done with a flow cytometry protocol adapted to phytoplankton, where single cells for the bottlenecks can be isolated by dilution after estimation of cell density using flow cytometry (46). To generate a clonal ancestral population (*T*_*0*_) for the MA experiment, the RCC623 culture density was estimated and a single cell was isolated by dilution and left to propagate for 2 weeks. Then, cell density was measured again, which allowed for the sampling of a volume corresponding to 10 cells, that was added to 10 mL of medium, of which 6 mL was used to inoculate 6 wells for each MA line on a 48-wells plate with 1 mL per well. Thus, each of the 6 wells statistically contains 1 cell. The 6 replicates limit the loss of MA lines due to technical errors or lethal mutations. Single-cell bottlenecks were performed every 14 days, by isolating 10 cells by dilution as described above from the first well with living cells encountered for each MA line; each bottleneck, 6 replicates from the same well were inoculated. Additionally, the number of generations *g* between single-cell bottlenecks can be calculated with *g=e(logN*_*t*_*/logN*_*0*_*)/t)-1*, with *N*_*0*_=1, *t*=14 and *N*_*t*_ as the cytometric count of the phytoplankton population. At the final time of the experiment, each MA line was transferred to 30 mL L1 medium and left to grow for 2 weeks in order to obtain enough cells for DNA extraction, yielding material for quality control, Illumina sequencing and PCR. DNA was extracted from 15 MA lines plus the ancestral line *T*_*0*_ using a modified protocol from (47) and (48): cell lysis was performed in two steps, first with Triton X-100 2% and in a second step with CTAB 4% and proteinase K 0.1 mg/mL.

### De novo mutation identification

Sequencing and library preparation were done at the DOE Joint Genome Institute (California, USA) using Illumina NovaSeq S4 to generate 150 pb paired-end reads (Table S13 for data accessions) that were mapped on the reference genome (14) with BWA mem2 v2.2.1 (49), after quality control with FastQC v0.12.1 (50). BAM files were sorted with samtools v1.20 (51). The callable genome (*G\**) was determined through 3 steps: i) sites require a minimal depth of 10 for all MA lines and 20 for the *T*_*0*_ (samtools depth with -Q20 -q20 options); ii) sites cannot have more than 2x the average depth, to exclude erroneous mapping in highly repetitive regions; iii) only the positions that met these criteria in all MA lines were kept in *G*. G\** analysis was done separately for the nucleomorph and the organelles, taking into account their relative ploidies (52), as well as scaffold 43 which appears to be duplicated in this strain (Fig. S14) and the 2 hypermutable regions. SNM and short ID calling was done with GATK Haplotypecaller v4.2.6.1 (53). VCF files containing all variants were then manually sorted to extract de novo mutations considering several criteria: i) candidate mutations are called in only 1 MA line, with up to 2 similar reads in other lines as sequencing errors; ii) at most 2 reads that do not support the candidate mutation; iii) the allele corresponding to the mutation is not present in the *T*_*0*_ genome; iv) the position is in *G\**. All insertions and deletions (ID) were manually checked in IGV (54). Then, primers were designed to verify all ID and a subset of SNM by PCR and Sanger sequencing (Table S14). All successfully sequenced amplicons (7 ID and 19 SNM) showed the expected mutation. The total mutation rate as well as mutation rates per type of mutation were calculated using the following formula:

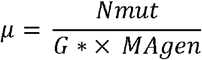

Where *N*_*mut*_ is the number of mutations, *G\** is the number of callable sites, and *MA*_*gen*_ is the sum of all independent generations across the 15 MA lines.

Additionally, we looked at the raw genomic depth with bedtools coverage v2-2.18.0 (55) using a window size of 1kb. This analysis provided evidence of large duplications in three MA lines (Fig. S2), as well as the duplication of scaffold 43 mentioned above.

To verify that the NCLDV hypermutable region is in one copy in the *B. natans* genome, we performed two analyses. First, we looked at the sequencing depth of these regions as compared to the nuclear genome using a window size of 1kb (Fig. S5); second we used Blastn to search the *B. natans* genome for sequences similar to the NCLDV sequence on scaffold 10 and found no similar sequences of even remotely the same size (biggest hit outside the region on scaffold 10 = 994bp on scaffold 111).

### Dinucleotide analysis

To identify sequences that might already have undergone TA→CA and TA→TG hypermutation we applied four different methods. First, we looked at the relative frequencies of these dinucleotides in genes previously identified as having transpoviron, virophage or NCLDV origins in (15). To get sequences, we extracted predicted CDS from the ‘Bigna1_filtered_transcripts.fasta’ on Phycocosm (56); this dataset contains a single spliced CDS (exons only) per gene model. We calculated the relative frequency of each dinucleotide, with reverse complements combined, using 1bp steps.

Second, we searched for sequences that are depleted in TpA or depleted in TpA and enriched in CpA+TpG in the whole coding genome. A subgroup depleted in TpA was extracted with Mclust in R v.4.4.3 (57) (Fig. S7). The 324 CDS identified were scanned for known Pfam domains (58) using HMMER hmmscan v3.2.1 (59) to see which domains were enriched in the TA-depleted subgroup; only hmmscan hits with E-value <10^-2^ were kept and each Pfam was counted once per sequence. Protein family and taxonomy were retrieved from InterPro (60).

Third, sequences of ≤300 bp in length were omitted to increase the representation of protein-coding CDS, resulting in 21,037 CDS. Relative dinucleotide frequencies were calculated as before to extract the 5% sequences lowest in TpA and the 5% highest in CpA+TpG; 470 sequences were shared. Hmmscan was run on these sequences with a stricter E-value <10^-5^. This time, each Pfam could figure multiple times per sequence for statistical analysis. Last, to extend the analysis to the whole genome, the genome was cut in 1kb windows and for each window, we estimated relative enrichment of both TpA and CpA+TpG dinucleotides using the Rho-value (18). This method uses nucleotide composition to predict expected dinucleotide composition within a sequence:

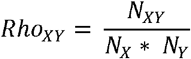

where *N*_*XY*_ is the raw count of any dinucleotide. A Rho >1 indicates enrichment and a Rho <1 depletion of the dinucleotide. Clusters depleted in TpA and enriched in CpA+TpG were defined as at least 3 subsequent windows with Rho_TpA_ ≤ 0.43 and Rho_CpA+TpG_ ≥ 1.52. Gene annotations were retrieved from Phycocosm (56).

Then, we compared the Rho_TpA_ of the NCLDV region of *B. natans* to its closest free living virus relatives. To determine the phylogenetic origin of the integrated NCLDV region, we used two complementary approaches. First, we searched the major capsid protein (aug1.10_g4662) of the *B. natans* NCLDV region on the NCBI refseq viral protein database using Blastp. The closest hit was *Heterosigma akashiwo* virus, a Phycodnavirus. To confirm that the NCDLV region could stem from Phycodnaviridae, we performed an independent annotation of the NCLDV region using getorf (https://www.bioinformatics.nl/cgi-bin/emboss/getorf, minimum size 300bp) to predict hypothetical proteins in the whole NCDLV region (sc10:336000-521500). The predicted proteins were searched against on the NCBI refseq viral protein database: 16 proteins had hits, and for 10 the hit with the highest bitscore was Phycodnaviridae. Thus, all complete Phycodnaviridae genomes (n=38, Fig. S12) were used for a RhoTpA analysis as described above.

### Single nucleotide polymorphism (SNP) analysis in the T_0_ strain

To determine whether we could detect signatures of the mutation rate spectrum we detected in the MA experiment over a longer time period, we analyzed the neutral SNPs in the viral hypermutable regions (NCLDV: scaffold_10:335999-521500, virophage: scaffold_146:16999-41997) that appeared between our *T*_*0*_ line (DNA extracted in 2021) and the reference genome (published in 2012) (14). Only intergenic positions with ≥20 depth (samtools depth, -Q20 -q20 options) in the *T*_*0*_ genome were considered. SNPs were called by GATK HaplotypeCaller and diversity was calculated as N_SNP_/N_total_, where N is the number of sites. All SNPs that showed a change from the reference genome from A/T to G/C in *T*_*0*_ were divided by the total number of A and T sites analyzed, giving SNP GC-rate. SNP AT-rate was calculated similarly and SNP AT-bias was calculated as AT-rate/GC-rate. TpA frequency was counted using the SeqinR count function (61) in R 4.4.3.

### Search for candidate deaminases involved in immunity

To investigate if *B. natans* possesses candidates for A-deaminases working on DNA, we aligned all 5 adenosine deaminases predicted in the *B. natans* genome (ProteinIDs: 142701, 73984, 135286, 57641, 78255) against a previously published alignment (21). The following sequences were retrieved from GenBank: ADAT2 from *Mus musculus* (MmADAT2, NP_080024.3), *Danio rerio* (DrADAT2, XP_005160675.1), *Branchiostoma japonicum* (BjADAT2, UDV79371.1), *Mizuhopecten yessoensis* (MyADAT2, XP_021373128.1), *Trypanosoma brucei* (TbADAT2, RHW71099.1), as well as their prokaryotic ortholog TadA from *Escherichia coli* (EcTadA, VWQ04824.1) and AID from *M. musculus* (MmAID, AAD41793.1). Protein sequences were aligned with the ClustalW algorithm (62) in MEGA11 (63), default parameters. Alignments were visualized with ESPript 3.0 (64) (File S1). AlphaFold predictions of Bn78255, BjADAT2 and TbADAT2 were done on AlphaFoldServer.com (65), using default parameters. Pymol v3.1.0 (66) was used for structural alignment of ‘model 0’ of each protein, default parameters (Fig. S13). To find a BjADAT3 homologue, its protein sequence was retrieved from GenBank (UDV79370.1) and was blasted against the *B. natans* genome with tblastn, minimum E-value <10^-2^, default parameters.

## Acknowledgments

We acknowledge the GenoToul Bioinformatics platform (Toulouse, France) for bioinformatics analysis support and cluster availability and all members and former members of the GENOPHY team, in particular Frédéric Sanchez and Anaïs Labécot for experimental procedures. We also thank the BIOPIC, BSBII and Bio2Mar platforms from the Observatory of Banyuls sur mer. We thank Max Emil Schön and Matthias Fischer (Max Planck Institute for Marine Microbiology, Bremen) for their help with the *B. natans* genome assembly and annotation. We thank Julien Henri for his help with the predictions and alignment of three-dimensional protein structures. The work (proposal: 10.46936/10.25585/60008631) conducted by the U.S. Department of Energy Joint Genome Institute (https://ror.org/04xm1d337), a DOE Office of Science User Facility, is supported by the Office of Science of the U.S. Department of Energy operated under Contract No. DE-AC02-05CH11231. This work was funded by the ANR ANR-23-CE20-0013 and ANR-21- CE02-0026.

