## Supplemental Figures for "Hypermutability of integrated sequences of viral origin in a Chlorarachniophyte"

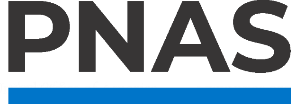

**Supporting Information for**

Ancient eukaryotic immunity through genome editing of viral sequences

Lisa Mettrop^1^, Anna Lipzen^2^, Gilles Mirambeau^1^, Kerrie Barry^2^, Igor V. Grigoriev^2,3^, Gwenaël Piganeau^1^, Marc Krasovec^1*^

^1^ CNRS, Microbial Biodiversity and Biotechnology Laboratory (UMR 8176), Sorbonne Université, Banyuls sur Mer, France.

^2^ DOE Joint Genome Institute, Lawrence Berkeley National Laboratory, California, USA.

^3^ Department of Plant and Microbial Biology, University of California Berkeley, Berkeley, California, USA

^*^**Corresponding author:** Marc Krasovec

**This PDF file includes:**

Figures S1 to S14

SI References

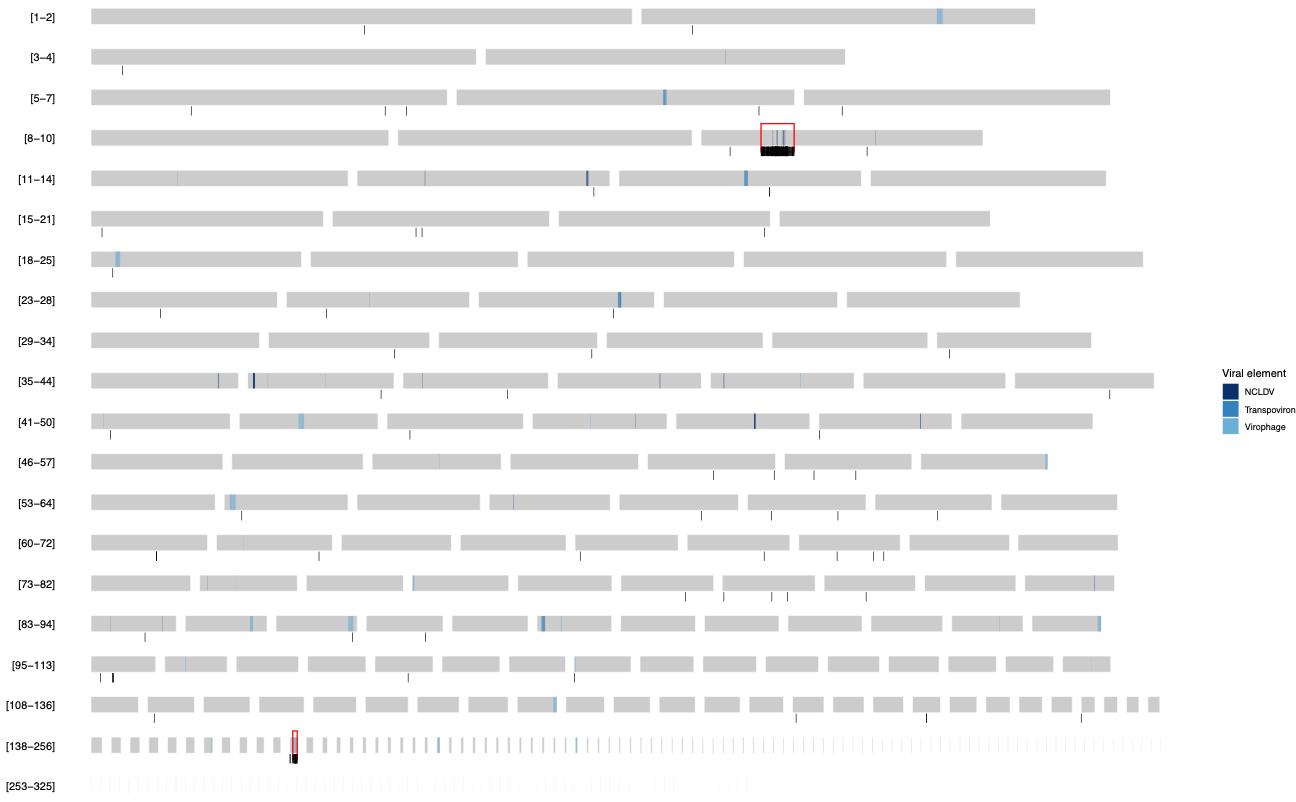

**Figure S1.** Genome architecture of *B. natans* in the assembly of (1) updated by the JGI. The nuclear genome is divided over 302 scaffolds. The number of the scaffolds on each row is indicated left of the graph; note that the numbering is not continuous, thus the last scaffold is called 325 instead of 302. Blue regions represent ORFs of different viral or transpoviron origins (2), which correspond to 0.41% (385,822 bp) of the total genome. Black vertical bars represent single nucleotide mutations and red rectangulars show the hypermutable regions exposed in our MA experiment.

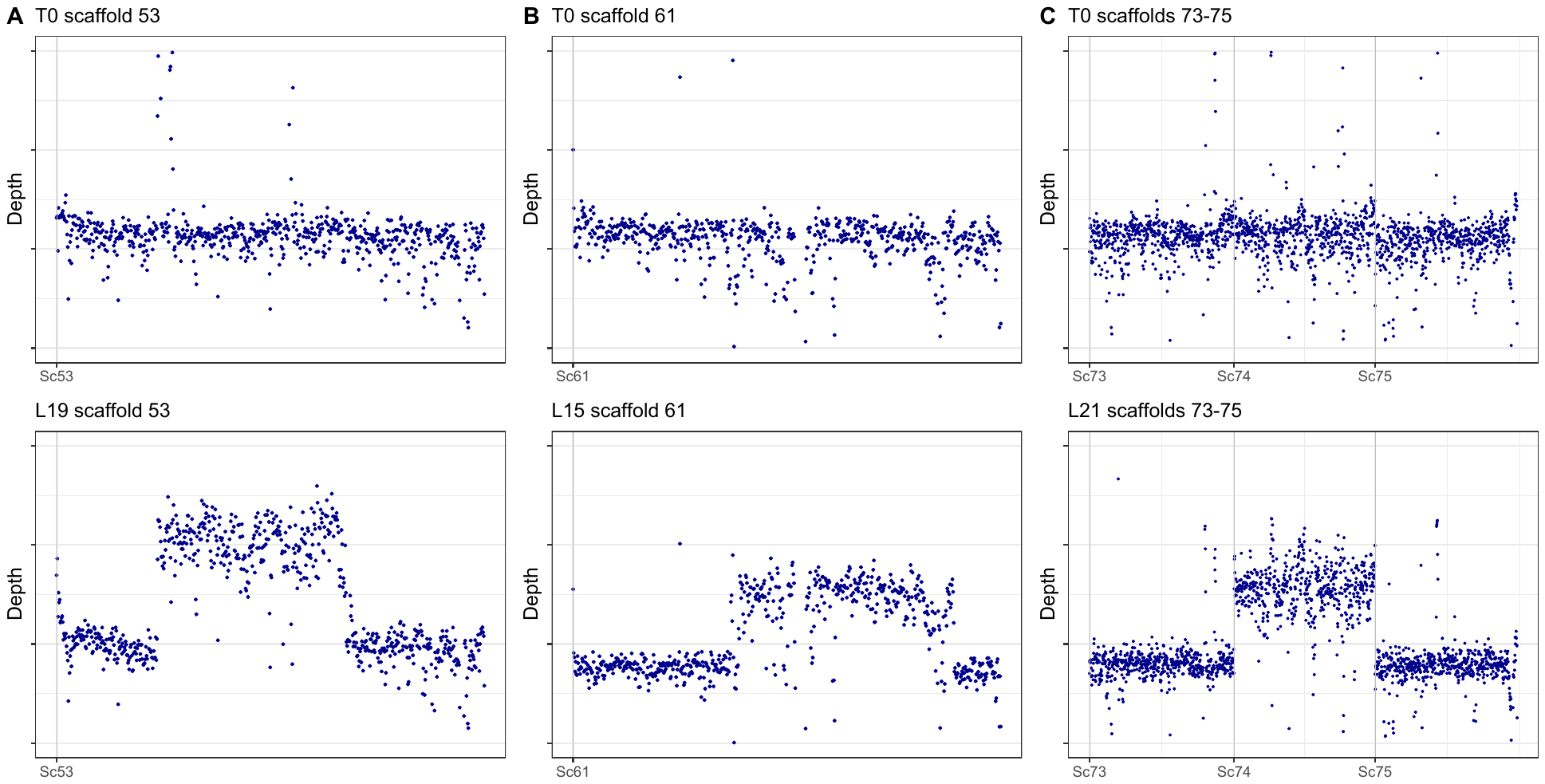

**Figure S2.** Raw genomic windowd depth showing large duplications in *B. natans* mutation accumulation (MA) lines. Genomic sequencing depth was calculated by 1 kb windows along the nuclear genome in the ancestral line (*T_0_* in the up panel) and MA lines in the bottom panel. Plots are zoomed on the scaffolds where three different MA lines present duplicated regions, visible as double sequencing depth. **A)** Line 19 shows a duplication of about 300kb in the middle of scaffold 53. **B)** Line 15 shows a duplication of about 332kb in the middle of scaffold 61. **C)** In line 21 the whole scaffold 74 is duplicated (about 544kb), scaffolds 73 and 75 are plotted around to show the difference in sequencing depth.

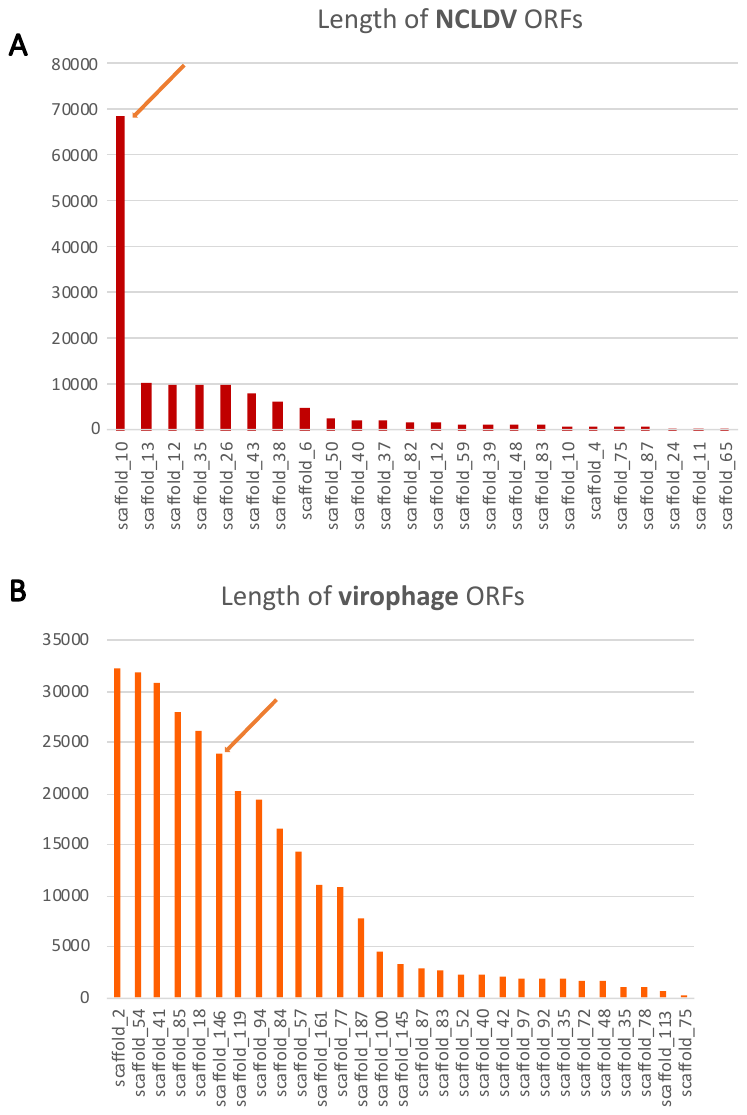

**Figure S3.** Length in bp of the viral elements identified in (2). The length of viral elements was estimated by combining the lengths of ORFs on the same scaffold that closely follow each other (<2000 bp between ORFs for virophages, <25 000 bp between ORFs for NCLDVs) plus the distances between combined ORFs. **A.** The hypermutable NCLDV region (showed by the orange) is the largest of identified integrated NCLDV elements in *B. natans.* **B.** The size of the virophage hypermutable region (showed by the orange) does not stand out from other virophage elements.

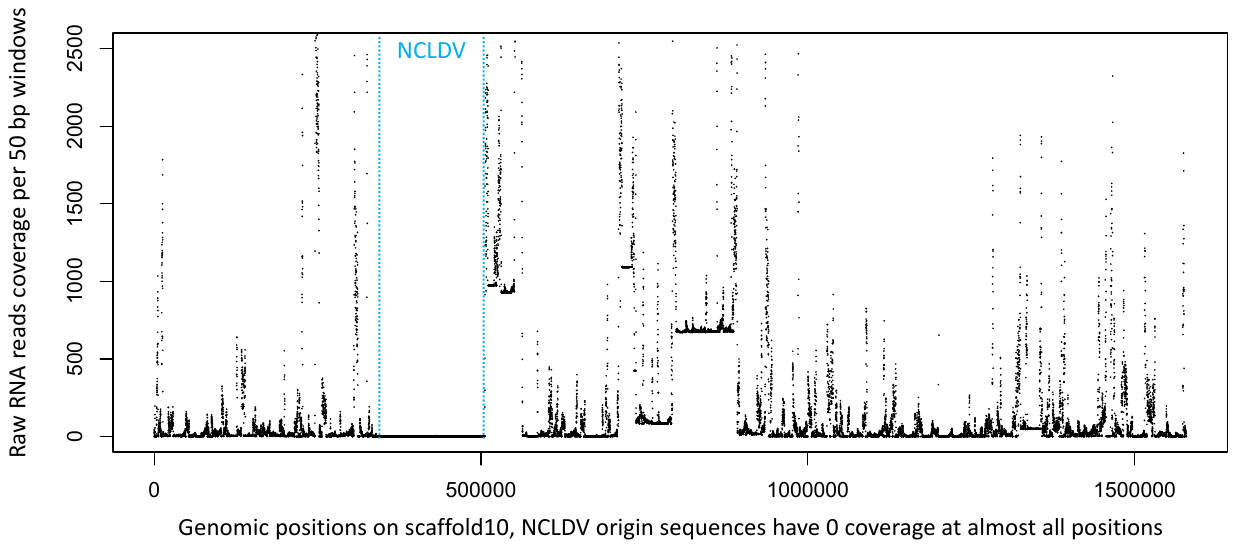

**Figure S4.** Raw RNA read depth on the scaffold 10 of *Bigelowiella natans*. RNA sequencing data comes from the MMETSP project (3), accession SRR1296871. Reads were mapped against the reference genome of *B. natans* (1, 2) with hisat2 and the bam files treated with samtools (4) and bedtools (5) to extract depth.

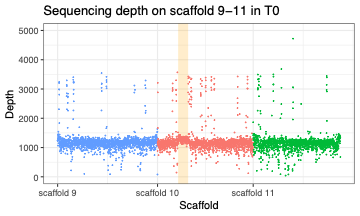

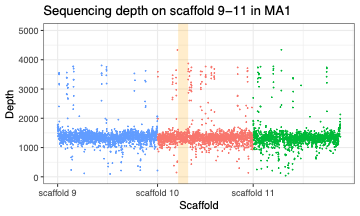

**Figure S5.** Raw windowed sequencing depth of 3 scaffolds (scaffolds 9 in blue, scaffold 10 in red, scaffold 11 in green) by 1kb windows from the *T_0_* of the mutation accumulation experiment of *B. natans* (upper panel) as well as MA line 1, the line with the highest mutation rate in the NCLDV region. The light orange bar indicates the hypermutable NCLDV region. On scaffold 10, 60/61 windows with a raw depth >2500 correspond to the same 5.7kb LTR TE, which is present in 150 copies in the *B. natans* reference genome, resulting in erroneous mapping and an inflated depth. The NCLDV region displays the average nuclear sequencing depth in both cases, confirning a single copy is present in the MA cultures as well as in the reference genome.

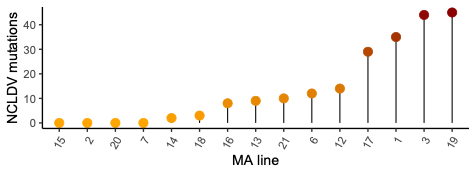

*µ_SNM_ = ~2 x 10^-6^*

45

**Figure S6.** The number of single nucleotide mutations in the hypermutable NCLDV region (y-axis) per MA line. (x-axis). MA line 3 had the highest mutation rate in this region, with *μ_SNM_* as high as 1.98x10^-6^.

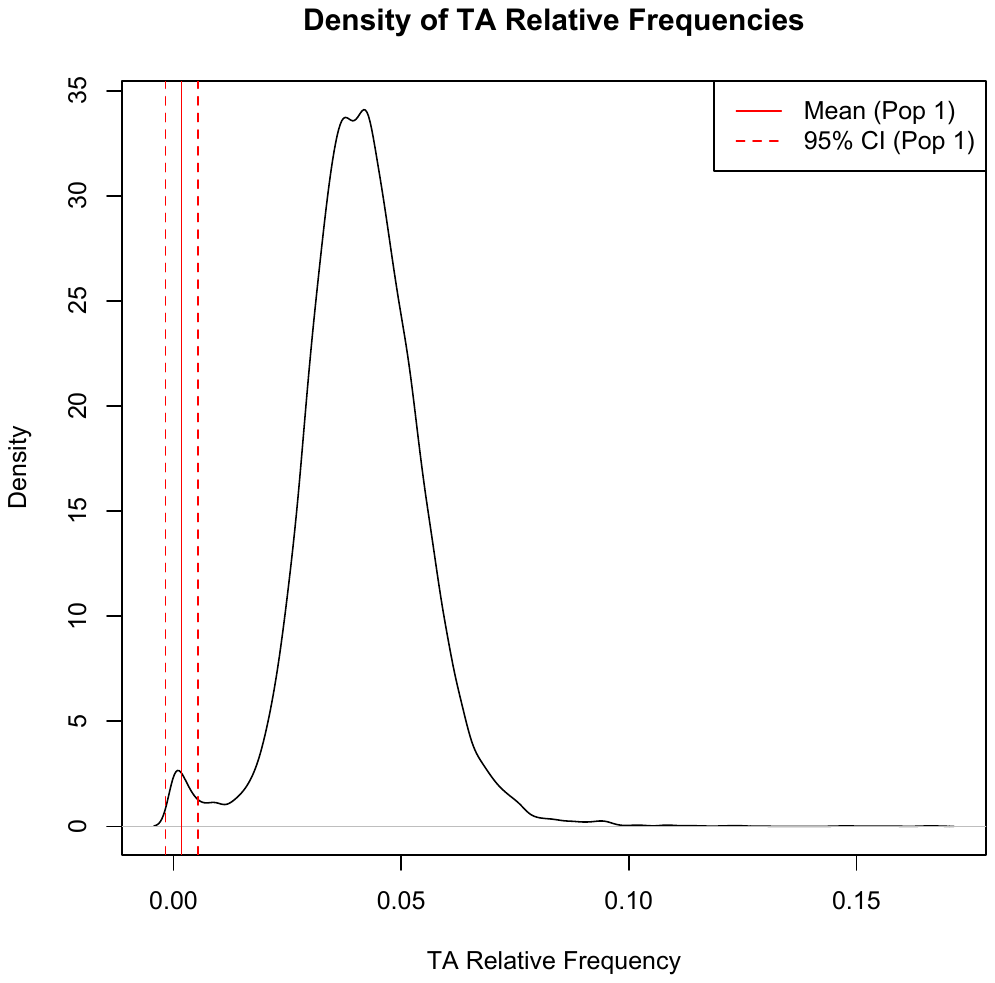

**Figure S7.** Distribution of relative frequency of TpA dinucleotides in *B. natans* predicted transcripts. Mclust (6) in R was used to extract the transcript subpopulation with extremely low TpA frequency; the mean frequency of this population (0.0018) is indicated with a full red line and 95% confidence interval with dashed red lines.

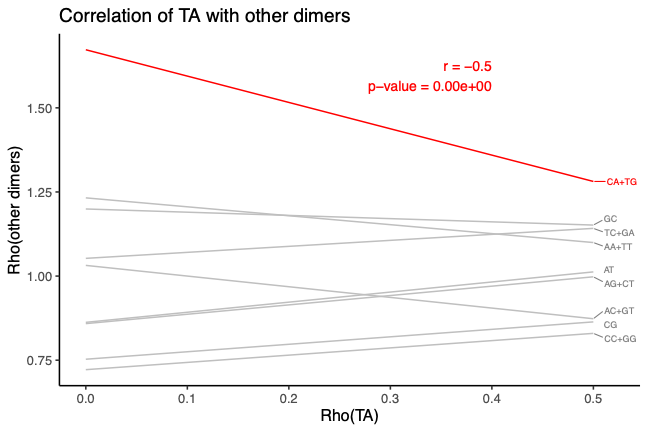

**Figure S8.** Pairwise relationships between Rho_TpA_ (7) and other dinucleotide Rho in the whole *B. natans* genome per 1kb genomic intervals but only considering windows with low TpA frequency (Rho_TpA_ <0.5). Red and grey lines represent linear regressions. The regression in red highlights the correlation between Rho_TpA_ and Rho_CpA+TpG_, the dinucleotide group with the strongest negative correlation to TpA; for this correlation, the Pearson correlation coefficient (r) with associated p-value is displayed in red.

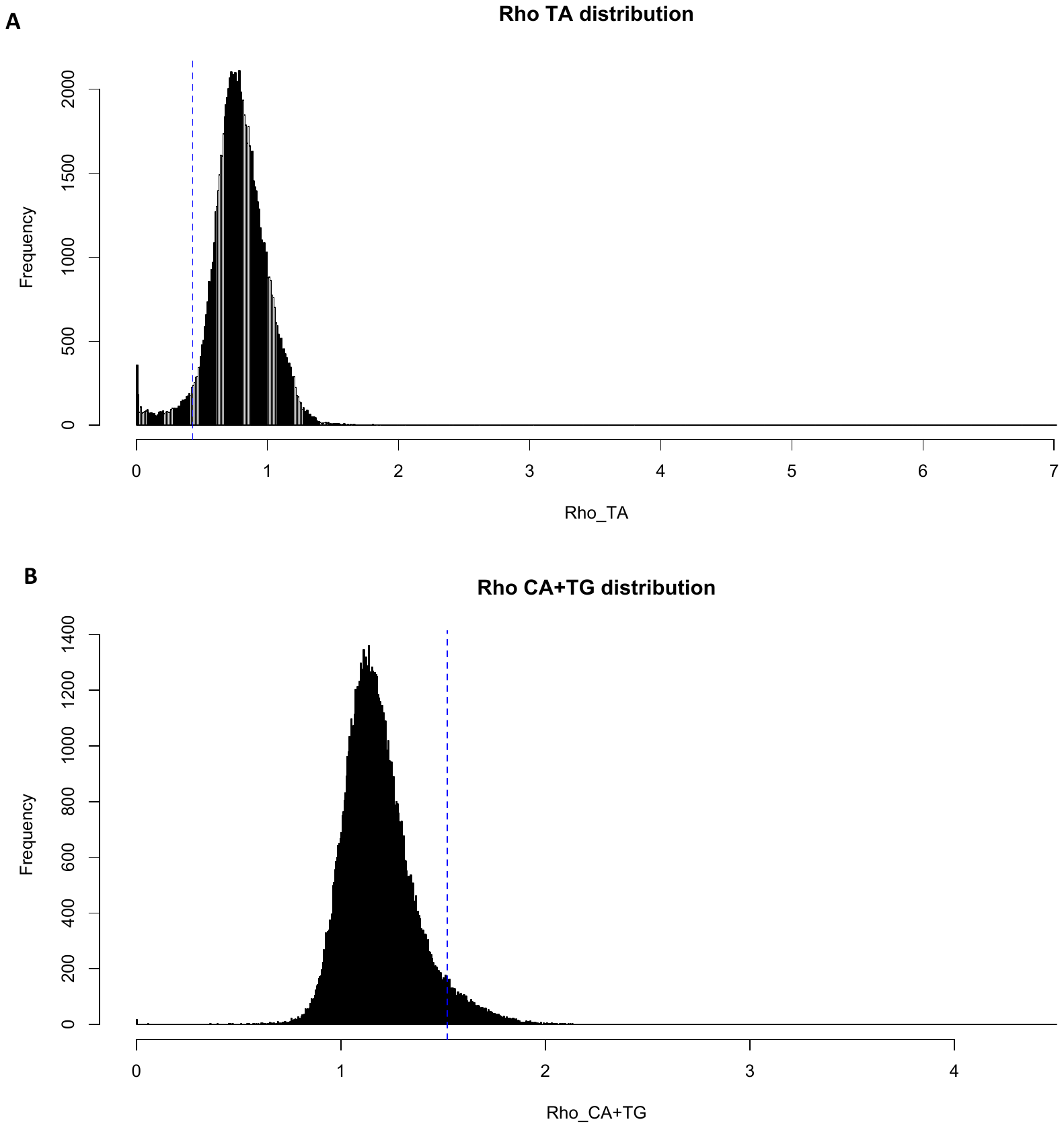

**Figure S9.** Distribution of Rho_TA_ (**A**) and Rho_CA+TG_ (**B**) in 1kb genomic windows. Dashed blue lines mark the 5% extremes used: 5% sequences with the lowest Rho_TA_ (≤ 0.43, upper panel) and 5% sequences with the highest Rho_CA+TG_ (≥ 1.52, lower panel). These cut-off values were used to extract the windows with the lowest TpA and highest CpA+TpG in oreder to analyse dinucleotide frequencies at a genome wide scale.

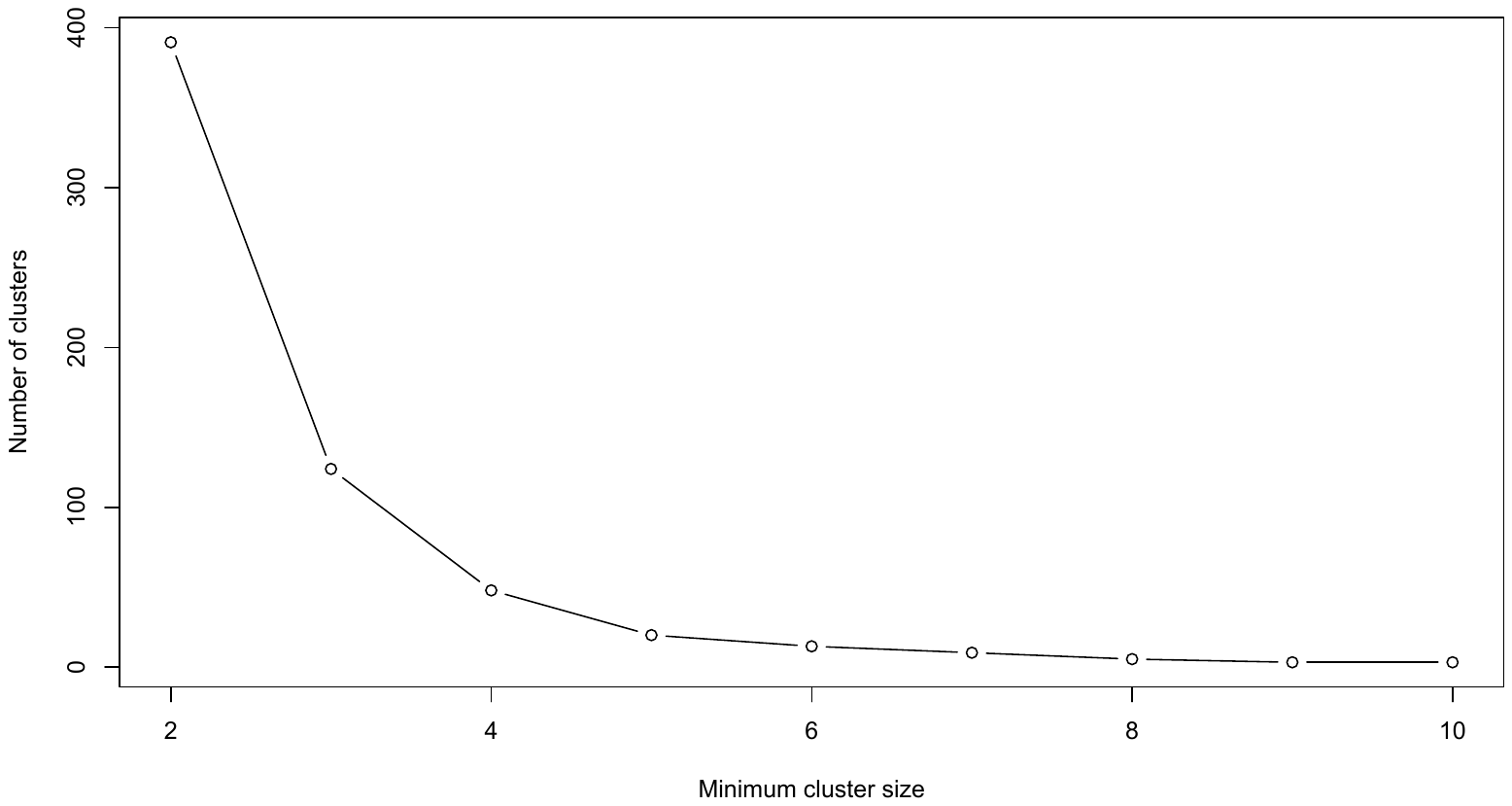

**Figure S10.** Distribution of the number of subsequent 1 kb windows depleted in TA and enriched in CA+TG, based on the Rho distributions in Figure S6. Following the ‘elbow rule’, clusters were defined as at least 3 subsequent windows with Rho_TA_ ≤ 0.43 and Rho_CA+TG_ ≥ 1.52.

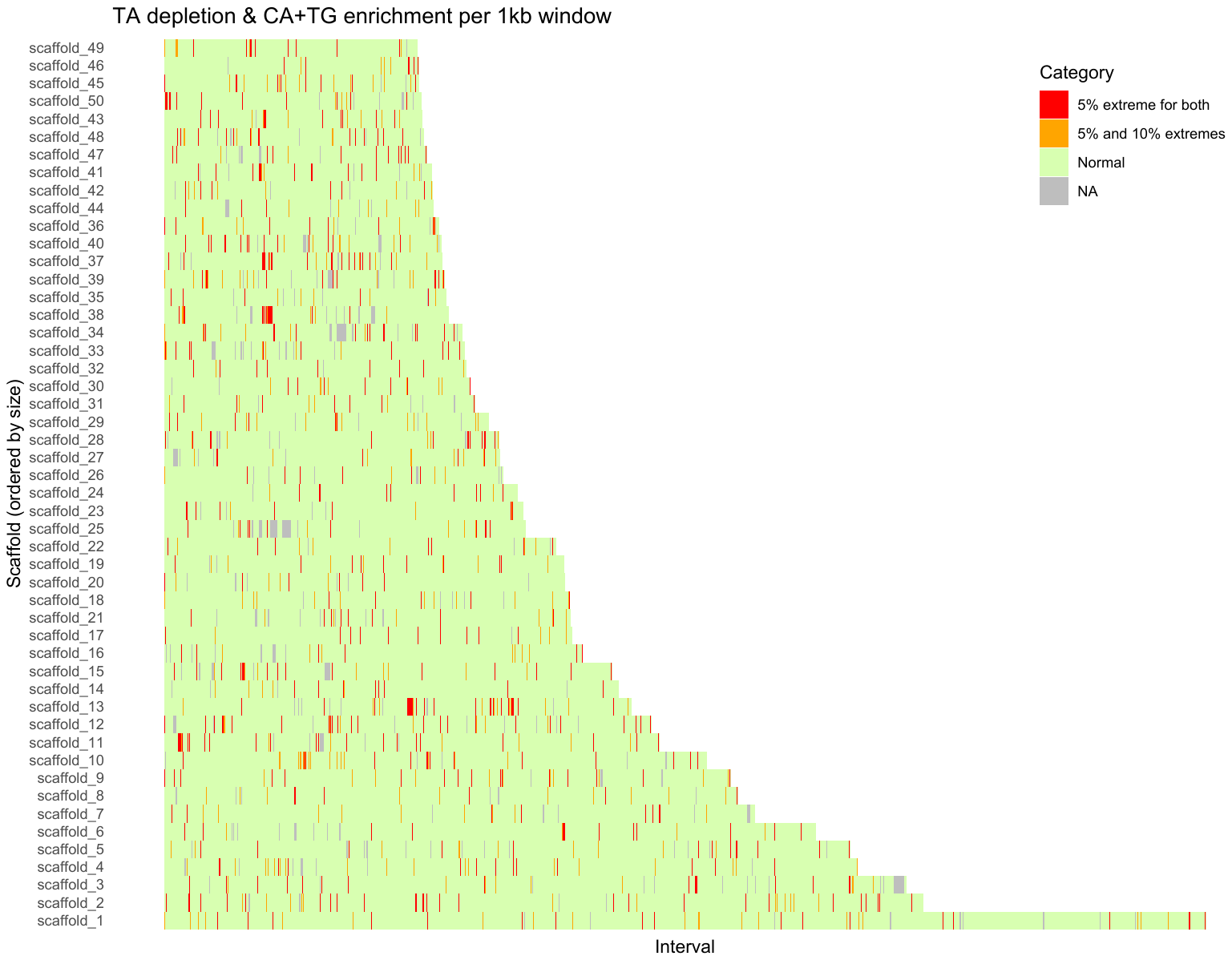

**Figure S11.** Distribution of Rho_TA_ and Rho_CA+TG_ values in 1kb windows across the *B. natans* 50 largest scaffolds. For each window, Rho_TA_ and Rho_CA+TG_ were calculated. In red, the windows that fall both in the lowest 5% Rho_TA_ (≤ 0.43) and highest 5% Rho_CA+TG_ (≥ 1.52). In orange, the windows that fall within aforementioned 5% extremes of either Rho_TA_ or Rho_CA+TG_ and in the 10% extreme of the other dinucleotide (Rho_TA_ <0.54 or Rho_CA+TG_ > 1.41). Grey corresponds to windows with missing or undefined Rho values and green to the windows with non-extreme values. This chromosomal painting highlights the spatial clustering of sequences both depleted in TA and enriched in CA+TG. A few clusters jump to the eye: scaffold_13 706,000-724,000, scaffold_35 285,000-288,000 and scaffold_50 1,000-19,000. These clusters are between 19 and 29kb in length and all contain multiple predicted genes (25 in total), however, only 2 genes have IPR predictions: on scaffold_13 there is a gene with proteinID 70820 that matches with IPR012337 (Polynucleotidyl transferase, Ribonuclease H fold) likely from a transposon, and on scaffold_38 there is proteinID 75972 that matches IPR001680, a WD40 repeat.

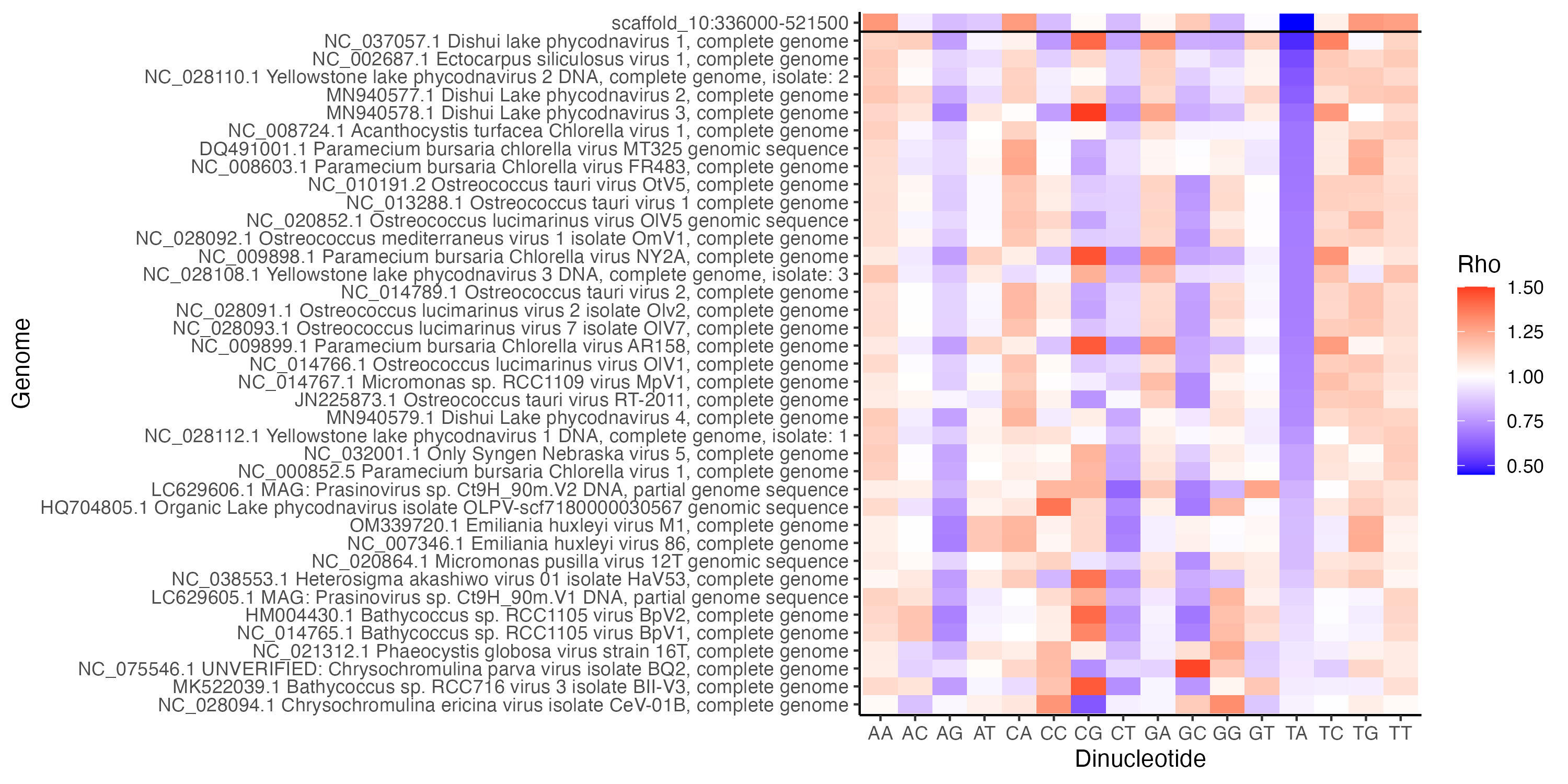

**Figure S12.** Dinucleotide Rho values for each genome of the 38 Phycodnavirus genomes available of the NCBI database including the NCLDV virus integrated in the *B. natans* genome (the first line on the top). The NCLDV integrated in the *B. natans* genome has the lowest TA frequency, in accordance with previous TA mutations such as observed in our experiment.

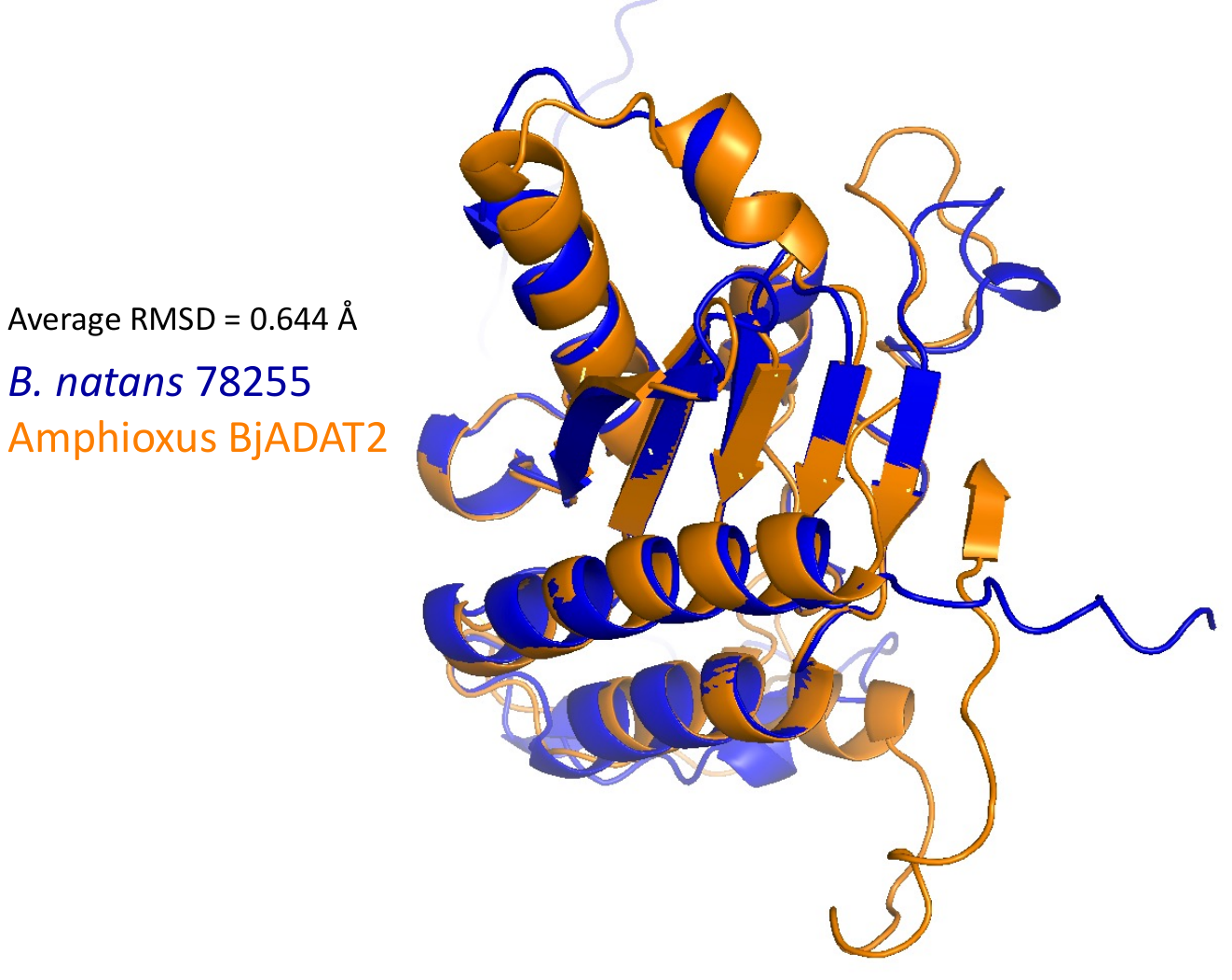

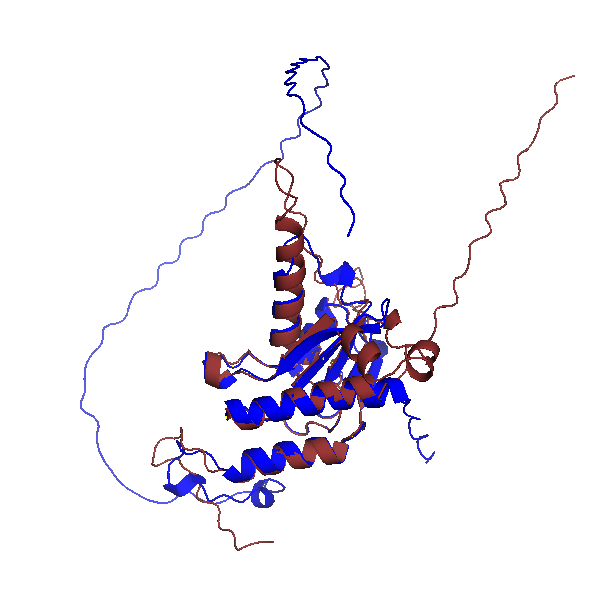

**Figure S13. Left:** Structural alignment of BjADAT2, the ADAR-like protein reported to target DNA in Amphioxus (8) (orange), and the ADAR-like candidate homolog from *B. natans* (blue). Protein structures were obtained from AlphaFold model 0 predictions (9) for Bn78255 and BjADAT2 and aligned in PyMOL (10). The resulting RMSD was 0.544 Å, indicating a highly similar structure. **Right**: Structural alignment of TbADAT2, the ADAR-like protein reported to target DNA in *Trypanosoma brucei* (11) (red), and the *B. natans* candidate homolog (blue). The alignment was performed as for the left panel and yielded an RMSD of 0.475 Å, meaning the structures are highly similar as well.

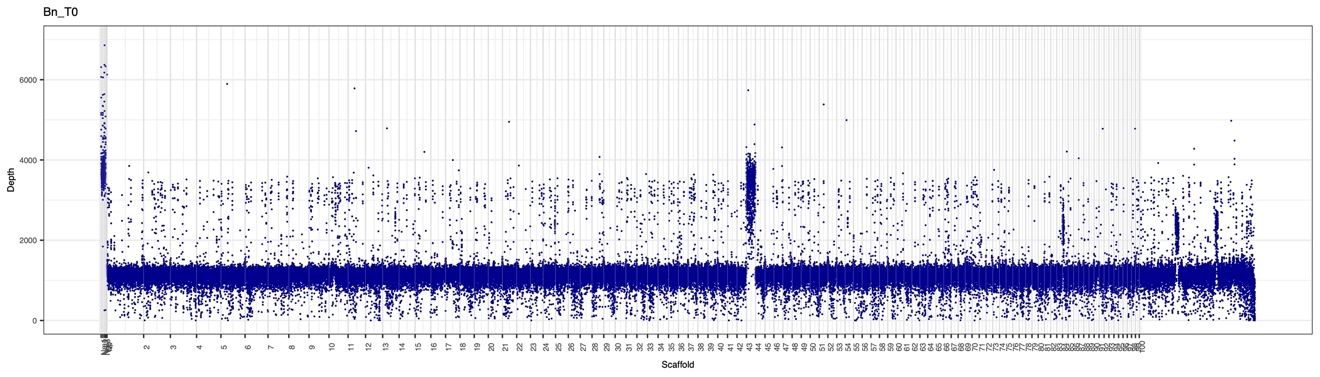
**Figure S14.** Illumina sequencing depth per 1000 bp window of the nuclear genome in the ancestral line (*T_0_*) shows the duplication of scaffold 43, visible as double sequencing depth.
