## Supplementary material for "Hypermutability of integrated sequences of viral origin in a Chlorarachniophyte": File S1

```
                  1                   10        20
BjADAT2     .....MADQGE.....................IEKYMYSALDLARAALAKGE........
BnADA78255  .....MQAGSRHN...............WTDRERKFMKMAIETARKAIVDEE........
MmAID       .....MDSLLMK.....................QKKFLYHFKNVRWAKGRHE........
MmADAT2     .....MEEKVESTTTPD......GPCVVSVQETEKWMEEAMRMAKEALENIE........
EcTadA      .....MSEVEFS....................HEYWMRHAMTLAKRAWDERE........
TbADAT2     ...MVQDTGKDTNLKGT......AEANESVVYCDVFMQAALKEATCALEEGE........
MyADAT2     .....MDEIHQ.......................KWMEKSFDYAEDALKSKE........
BnADA57641  .....MSEQVRDKVILT....VEIKSLNASILPQKKVENCYIGKDRVIQVTDAFIR....
BnADA73984  .....MVQTALFLSLLG........LALSFDPSTSNILQNRPRVELHIHLDG........
BnADA135286 .MASSVDDATRSPILAQKKEAALSPSLVSSSSTTSFAETVASLVMDAYDKLERRGKPQRQ
BnADA142701 MWTAIVVDLLVVLVCIS......HYVYFPMIFQTTKLGSVRDETLLELMKDDIG......

                   30                           40        50        60
BjADAT2     ...VPVGCVMVYQ..................GQIVAQGHNRVNESKNATVHAEMVAVGQL
BnADA78255  ...VAVGCVFVDTRT...............DRVIAATHNETTKTRNASRHCELVAVDQI
MmAID       ....TYLCYVVKRR.................DSATSCSLDFGHLRNKSGCHVELLFLRYI
MmADAT2     ...VPVGCLMVYN..................NEVVGKGRNEVNQTKNATRHAEMVAIDQV
EcTadA      ...VPVGAVLVHN..................NRVIGEGWNRPIGRHDPTAHAEIMALRQG
TbADAT2     ...VPVGCVLVKADSSTAAQAQAGDDLTLQ.KLIVARGRNATNRKGHALAHAEFVAVEEL
MyADAT2     ...VPVGCILVYRG.................EEVIGVGRNEVNETKNASRHAEIVAIDQV
BnADA57641  ..DIPKTDLHLHLDG....SVRLETLIELAKQQGVELPASTPEGLREKVFKDTYKDLPEY
BnADA73984  ..SVPVPSLFAIAT.................EKEVHIPGIGIPKSTADIWRALDNVTSSW
BnADA135286 EWTVLAGIIRTHECGESGSILSSDVISIATGTKCVGRKNINPNGETLNDLHAESLVRRAF
BnADA142701 ...ATMAELEEMRVQLNDKRTLSQCFKIFGLIHKVVKTHAAITRITVEVLEDFSNDGVRY

                 70
BjADAT2     LEWCMKVGLD..................................................
BnADA78255  LG..RDASLR..................................................
MmAID       SDWDLDPGRCYRVT..............................................
MmADAT2     LDWCHQHGQS..................................................
EcTadA      GLVMQNYRLI..................................................
TbADAT2     LRQATAGTSENIGGG.............................................
MyADAT2     LSWCVHRDMN..................................................
BnADA57641  LQGFQYTTAVMRIGSAMERIAYELGVDCFKEGIRYIEVRFAPQLHIDPSAKKSGGDGKED
BnADA73984  QRFDVVNNIIGG................................................
BnADA135286 VRYLLEDIRKAAHRGGASQSENFVKATQVENIDSRGETAYASPPPPPPPPPQSPPSPSSK
BnADA142701 VELRTTPRIDAKHN..............................................

                         80        90
BjADAT2     ..........KNEVLPIVQLYVTCEPC.................................
BnADA78255  ..........PEDIFPHCELYVTVEPC.................................
MmAID       .......WFTSWSPCYDCARHVAEFLR.................................
MmADAT2     ..........PSTVFEHTVLYVTVEPC.................................
EcTadA      ................DATLYVTLEPC.................................
TbADAT2     .....GNCGAVSQDLADYVLYVVVEPC.................................
MyADAT2     ..........PKVVFGQSVLYVTVEPC.................................
BnADA57641  GDANIDKENNPLSDCVSAIYYVSKGLKRASEEWNSREEVKDGDEPGYG............
BnADA73984  .........DPDSLASVAEDFVARQANQG...............................
BnADA135286 DGLNKRQCYYRIADGIRFHLFISQPPCGDASIYHRTSISKTGGNSEERKELGIGNKRKKP
BnADA142701 .........MTQESYVEAVLKAATSILLNEFHSLEQDFTS....................

BjADAT2     ............................................................
BnADA78255  ............................................................
MmAID       ............................................................
MmADAT2     ............................................................
EcTadA      ............................................................
TbADAT2     ............................................................
MyADAT2     ............................................................
BnADA57641  ............................................................
BnADA73984  ............................................................
BnADA135286 DSREGIVDEPSSSSSQQQQEQRQQYHQQQQQQHPNLNQTGAKILLDDDSINLRRTVRRVG
BnADA142701 ............................................................

                                                          100
BjADAT2     .........................................IMCAGALRLLNIP......
BnADA78255  .........................................IMCAAALRIIKIG......
MmAID       ...........................................WNPNLSLRIFT......
MmADAT2     .........................................IMCAAALRLMKIP......
EcTadA      .........................................VMCAGAMIHSRIG......
TbADAT2     .........................................IMCAAMLLYNRVR......
MyADAT2     .........................................VMCAGALRQVGVP......
BnADA57641  ......................................YGIICCAMRMIYKQLSPYYRSL
BnADA73984  ..........................................VGYTEVRYDPVR......
BnADA135286 REDVQRQLIGVPRLKSGRSDIPKELRTLSMSCSDKIAKWLFCGCQGGLLANLITGPIKIS
BnADA142701 .........................................LRIEVEIAHKTVA......
```

```
                                                   110          120
BjADAT2       ....................................VVWYGCPNPRFGGCGSVLSVHSDR
BnADA78255    ....................................RVVYGCANPRFGGTGSVLFLHDTK
MmAID         ....................................ARLYFCEDRKAEPEGLRRLHRAGV
MmADAT2       ....................................LVVYGCQNERFGGCGSVLNIASAD
EcTadA        ....................................RVVFGARDAKTGAAGSLMDVLHHP
TbADAT2       ....................................KVYFGCTNPRFGGNGTVLSVHNSY
MyADAT2       ....................................LVVYGCANDRFGGCGSVLSVNEDE
BnADA57641    CALHEHEPMRKKVAYASRALVASAVFARDELGLPVVGIDMAGAEKGFPAEWNMDAFNYAK
BnADA73984    ....................................MAKSMYTGKSISEEDAIKAISGGL
BnADA135286   TIVVSKPHNASVEVKKSGLDSSSSSSS........SSSNYGSRSTRGKSCSSTSSLESKK
BnADA142701   ...................................LAKAYQTCKDR.RNQGVVVGVDFSG

                 130                   140            150          160           170
BjADAT2       LESYGQ............QFKCIGGIFAEPC...AALLKQFYGQQNPNAPNPKK.KSEGR
BnADA78255    R.SEGL............GYECIGGLDEQQA...IALLKSFYAKGNPNAPAAKR.ARSLE
MmAID         QIGIMTFK.........DYFYCWNTFVENRE....RTFKAWEGLHENSVRLTRQLRRILL
MmADAT2       LPNTGR............PFQCIPGYRAEEA...VELLKTFYKQENPNAPKSKV.RKKDC
EcTadA        GMNHRV..............EITEGILADEC...AALLSDFFRMRRQEIKAQKK.AQSST
TbADAT2       KGCSGEDAAI.......VGYESCGGYRAEEA...VVLLQQFYRRENTNAPLGKR.KRKDL
MyADAT2       LPTLGP............SFQCISGVMGDKS...VVMLKDFYKGENSNAPDGKR.KIKNT
BnADA57641    KNFMNKTVHAGEGYGPGSIFQAITDLHASRIGHGFHLFSWDLAMQQSQSGKTQK.EAKQY
BnADA73984    RKGAETHG........VAVYSLLCAMRGTNA...SSCDKECFRKAKEQLGLNTT.VHAGE
BnADA135286   KKKKDEKIST.......PPRLLICNRQYPRS...LANIRNDYHSHSKETSRKNK.RIRSE
BnADA142701   NPHAGSFAQFR......PAFEAAKKAGLRCAIHCAEIEAEDEVEDILRFGADRLGHAIVM

                    180                   190        200
BjADAT2       STGLGGNLASNFG........SAFGGSGDGSEYYAAGYGYTEF.................
BnADA78255    SRGGGGGGGDDNDDATLDPPSSSVPPSGNVSHNDKAKVAHGKEKRGKDNEEEKNCSNTGK
MmAID         PLYEVDDLRDAFRMLGF...........................................
MmADAT2       Q...........................................................
EcTadA        D...........................................................
TbADAT2       SVV.........................................................
MyADAT2       D...........................................................
BnADA57641    VSSLVEWVCDKRITLEVCLTSNLQTMPELKGNVNNHVFGKMVDRQLSVTLCTDNRLVSNT
BnADA73984    AVRPHDVLTAVRDMKVDRIGHGYAATEDAEALKALKDYGVHLEACPPGANRHHVIDAIGV
BnADA135286   SQHDKEQIKRRKGEMGRKVSNDIRRSPSACGLALNWYKGGAVEVLGGTTGRKQGANMRQK
BnADA142701   STNHKELLSKMGTPVEICPTSNMRTLQILDIKQHP.TLGLWLQNGHPFSICTDDFGVFRT

BjADAT2       ............................................................
BnADA78255    RKGFDERSCDD.................................................
MmAID         ............................................................
MmADAT2       ............................................................
EcTadA        ............................................................
TbADAT2       ............................................................
MyADAT2       ............................................................
BnADA57641    TIPKEYRLAIDNFGLTLKQLRDITICGFKRSFSPEPYKQKREYVRKAINYFDAMQPCVSK
BnADA73984    YKEMKLSFGLNEDDPTVYFENCTMDSVEALTREHLSFTDADMRKAYADAFAARFGPHTPK
BnADA135286   SIKTCSRLCKAALWTLFRDVYAGINGKSVFSSYFEAKSCSRWYRELKSDFKLSMRQWVRK
BnADA142701   SLSEETRLVATAFGLSLKQVADISLRALEMAFIPDKDELCRLRKRFETEINAILISQEKH

BjADAT2       ......................
BnADA78255    ......................
MmAID         ......................
MmADAT2       ......................
EcTadA        ......................
TbADAT2       ......................
MyADAT2       ......................
BnADA57641    NIKF..................
BnADA73984    ......................
BnADA135286   ERADKISILSSHQFQQVNRNIN
BnADA142701   EI....................
```
